# Extracellular Matrix protein gelatin provides higher expansion, reduces size heterogeneity, and maintains cell stiffness in a long-term culture of mesenchymal stem cells

**DOI:** 10.1101/2021.12.20.473506

**Authors:** Pankaj Mogha, Shruti Iyer, Abhijit Majumder

## Abstract

Extracellular matrices (ECM) present in our tissues play a significant role in maintaining tissue homeostasis through various physical and chemical cues such as topology, stiffness, and secretion of biochemicals. They are known to influence the behaviour of resident stem cells. It is also known that ECM type and coating density on cell culture plates strongly influence *in vitro* cellular behaviour. However, the influence of ECM protein coating on long term mesenchymal stem cell expansion has not been studied yet. To address this gap, we cultured bone-marrow derived hMSCs for multiple passages on the tissue culture plastic plates coated with 25 μg/ml of various ECM proteins. We found that cells on plates coated with ECM proteins had much higher proliferation compared to the regular tissue culture plates. Further, gelatin coated plates helped the cells to grow faster compared to collagen, fibronectin, and laminin coated plates. Additionally, the use of Gelatin showed less size heterogeneity among the cells when expanded from passages P3 to P9. Gelatin also helped in maintaining cellular stiffness which was not observed across other ECM proteins. In summary, in this research, we have shown that gelatin which is the least expensive compared to other ECM proteins provides a better platform for mesenchymal stem cell expansion.

## Introduction

The extracellular matrix (ECM) proteins such as collagen, elastin, fibronectin, and laminin form the scaffold of tissues on which the cells anchor. ECM proteins play important roles in proper functioning of tissues by providing the necessary physical and chemical cues. During *in vitro* cell culture, the plastic plates are often coated with a suitable ECM protein for better cell adhesion and cell survival [1]. It has been demonstrated that cells adhere to different ECM proteins with affinities in the following order, fibronectin > collagen I ≥ collagen IV ≥ vitronectin > laminin-1 through different integrins [1]. The effect of ECM proteins on various cellular responses such as migration, adhesion, proliferation, differentiation, have been extensively studied with different cells, including stem cells [1–3]. Peptides mimicking ECM proteins have also been shown to influence cellular behavior *in vitro* [4]. However, the reported studies investigating the effect of ECM proteins on cellular behaviour have been done for a small culture duration, usually 48-72 hours. The effect of ECM protein-coated substrates on long term culture or expansion of stem cells has not been investigated.

In this work, we aimed to study the effect of ECM protein-coated plastic on long-term expansion of bone marrow-derived human mesenchymal stem cells (BM-hMSCs). While BM-hMSCs have huge therapeutic potential, one of the major bottlenecks is to have enough number of cells required for cell therapy and regenerative medicine [5,6]. As the availability of hMSCs from bone marrow is low, *in vitro* expansion is the only possible way to generate large number of hMSCs. However, after a definite number of cell divisions, all primary cells enter a permanent growth arrest known as replicative senescence which limits their expansion and differentiation ability. Hence, methods are to be explored that may generate a greater number of cells starting from a relatively small number before they enter replicative senescence. Earlier, our group has shown that using collagen coated optimally soft Polyacrylamide gel as substrates, the replicative senescence can be delayed in hMSCs and hence a greater number of cells can be generated [7]. In this work, we explored the effect of ECM on hMSC growth rate. We coated the plastic cell culture plates with ECM proteins namely, Gelatin, Collagen, Fibronectin, and Laminin, and investigated their effect on cell morphology, cell stiffness, doubling rate, and final cumulative population doubling till growth arrest. The result shows that coating the plastic plates with any ECM improves the doubling rate and final cell count but does not have any effect on the duration of culture till the growth arrest. Among all the different ECMs used, gelatin was found to be the most effective in increasing cell proliferation. As gelatin is co-incidentally also the cheapest ECM among all the ECM tested, our work suggests an inexpensive and simple modification in the culture process to have higher number of viable stem cells from *in vitro* expansion.

## Material and method

### Cell culture

Primary bone marrow hMSCs (BM-hMSCs) were obtained from Lonza (PT-2501) at passage 2. The cells were expanded and used from P3 onwards for experiments. Cells were cultured in low glucose DMEM (Himedia Al006) supplemented with 16% FBS (Himedia RM9955/ Gibco 12662029), 1% glutamax (Gibco 35050061) and 1% penicillin–streptomycin (Gibco 15140122), maintained at 37°C with 5% CO_2_. Upon reaching confluency, cells were trypsinized using TrypLE express (Gibco 12604013).

### ECM protein coating and cell seeding

For coating the ECM protein onto T-25 flasks (Nunc 156367 tissue culture flask with 25 cm^2^ culture area), 3 ml of Dulbecco’s Phosphate Buffered Saline (DPBS) (Himedia TS 1006) containing ECM proteins at a concentration of 25 μg/ml was added to the flask and incubated at 4°C overnight. For example, a T25 was incubated with 3 ml of a 25 μg/ml collagen1 (Gibco A1048301) solution made in DPBS. Similarly, for laminin (Sigma L2020), fibronectin (Sigma F4759), and gelatin (Sigma 1393), 3 ml of respective ECM protein solutions (25 μg/ml) were added in respective T-25 flasks and incubated overnight at 4°C. For a non-ECM negative control, the T-25 flask with a proprietary coating by the manufacturer was incubated with 3 ml of DPBS only. Before seeding the cells, each ECM protein-coated TCP was liberally washed with DPBS thrice and acclimatized to room temperature.

### PD and CPD calculation

After four hours of cell seeding, at least 15 images were acquired from each flask sampling the entire culture area. The population doubling for each passage were calculated as previously described [7,8]. Briefly, total number of cells per frame were manually counted and divided by the total frame area (cm^2^) to get the seeding density. The average seeding density was then multiplied by the total area of the flask (25 cm^2^) to obtain the total number of cells seeded for each condition. Before trypsinization, random images were again captured and total cell number was obtained as explained above. Using the following equations, population doubling (PD) and cumulative population doubling (CPD) were calculated:

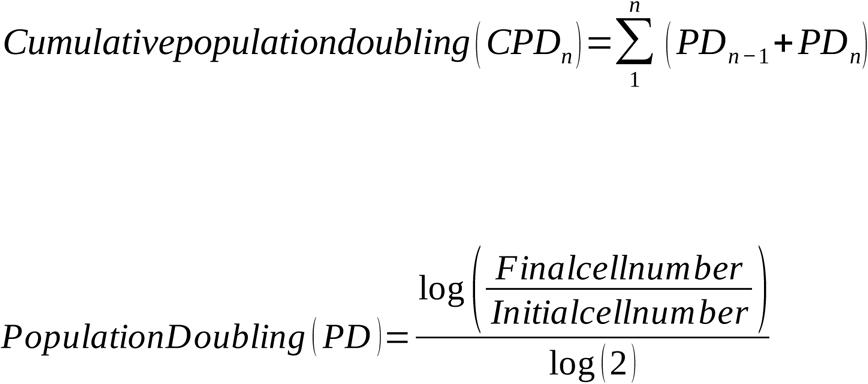

Where, ‘n’ denotes the number of passages for which the calculation was done.

### Cell spread area quantification

The images captured before trypsinization were used to calculate the cell spread area by tracing the cell’s periphery using ImageJ after applying an appropriate scale bar.

### Atomic force microscopy (AFM) stiffness quantification

Atomic force microscopy measurements were done with a PT.PS.SI.4.5 (Novascan) probe attached with a polystyrene bead of 4.5 μm diameter on MFP-3D (Asylum Research) under contact mode force. The spring constant and frequency ranged from 0.09-0.27 N/m and 26 to 28 kHz respectively. The force curves were obtained at the regions away from the cell nucleus and the cell periphery. The force curves were then fitted with hertz model provided within the software from Asylum research after setting the correct parameters for tip geometry, material of the probe, and Poisson’s ratio of the material. The stiffness values were recorded for indentation up to first 500 nm from the contact point.

### MSC differentiation

Cells from early passage P4 and the cells of passage P8 from gelatin coated TCP and uncoated TCP were seeded onto a 12 well plate at a seeding density of 10000 cells/cm^2^. Upon reaching confluency, the cells were switched from MSC culture media to differentiation media Adipogenic media (Gibco-12662029), and Osteogenic media (Gibco-A1007201)). Differentiation media change was given every second day for next 14 days for Adipogenic differentiation and 21 days for Osteogenic differentiation. After the completion of differentiation duration, cells were fixed with 4% PFA for 20 min at room temperature followed by staining with Oil Red O (Sigma O0625) for adipo differentiation cells and Alizarin Red S (Sigma A5533) for osteogenic differentiation as explained earlier [9]. The cells were incubated with staining solution for 20 mins and then washed thrice with DPBS (adipo) or MilliQ water (osteo), the cells were then imaged with Evos FL auto (ThermoFisher scientific) in bright-field colour channel.

## Results

### ECM protein coating enhances the proliferation rate of mesenchymal stem cells *in vitro*

To investigate the effect of ECM protein coating on cell proliferation, BM-hMSCs were sequentially cultured on commercially available plastic tissue culture plates (TCP) by coating them with the ECM proteins and compared against a commercially available (Nunc Gibco) non-ECM coated TCP flask as a control (figure 1A). A complete media change was done on every 3^rd^ day and the culture was passaged on every 6^th^ day. After trypsinization, cells were reseeded on freshly ECM protein-coated plates or TCP at a density of 1000 cells/cm^2^. The process was continued till there was no increase in cell number indicating growth arrest.

**Figure 1:**
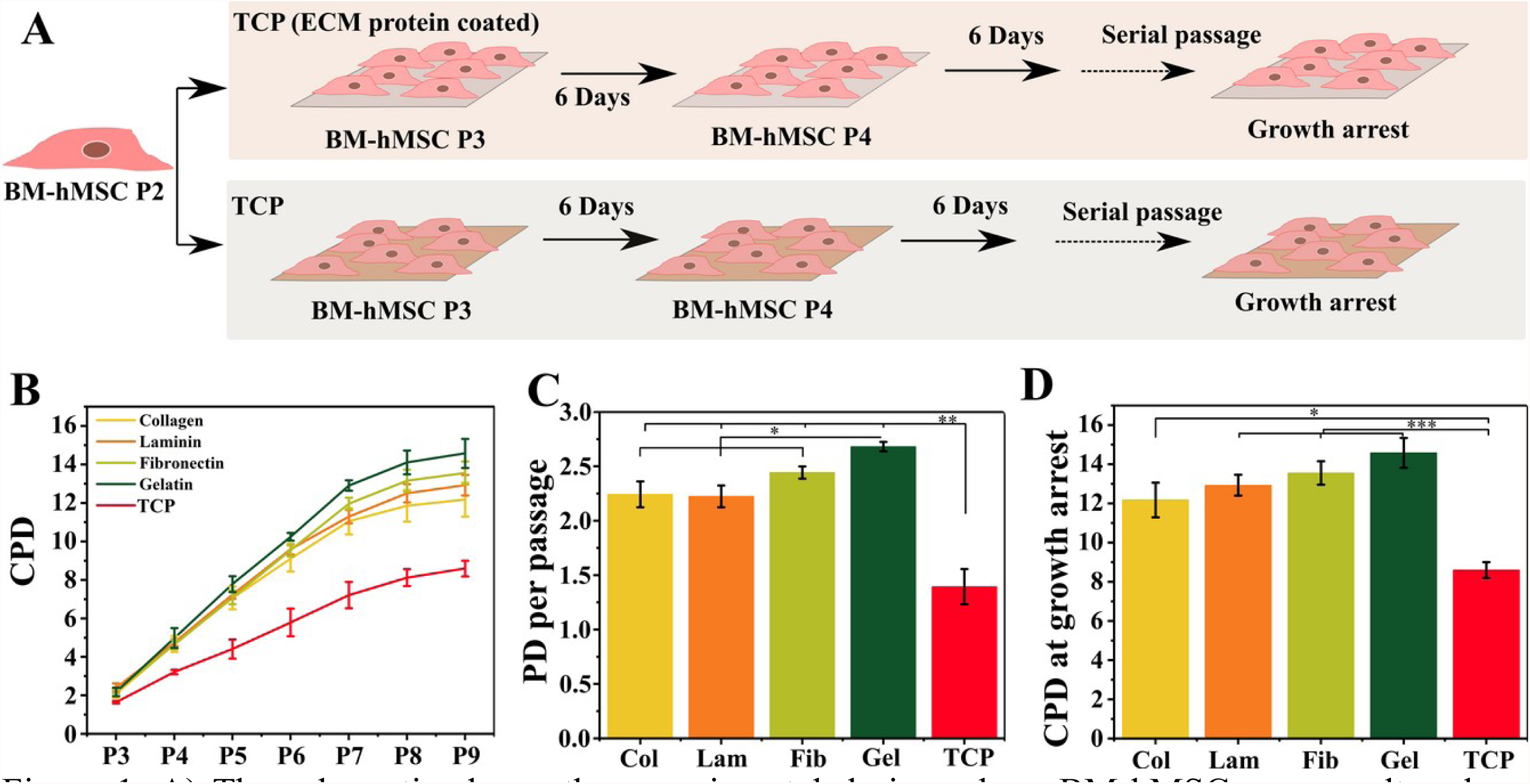
A) The schematic shows the experimental design where BM-hMSCs were cultured on tissue culture plates, either coated with or without ECM-protein. The cells were routinely cultured on respective TCP flasks until they reached growth arrest. B) Line graph shows the cumulative population doubling of the cells cultured on different coating on TCP. C) Bar graph shows the population doubling per passage till P7. D) Bar graph shows the final cumulative population doubling values obtained at cell growth arrest for each coating. *P-*value *=p<0.05, **=p<0.01, ***p<0.001.

First, we assessed the viability of BM-hMSCs on ECM coated and non-coated plates through Calcein and Propidium Iodide (PI) staining. The cells showed similar viability across all ECM protein-coated and non-coated TCP as shown in figure S1. Next, we calculated the growth by calculating cumulative population doubling (CPD) as explained in the methods section and plotted the same against the passage number as shown in figure 1B and figure S2. Results derived from this data show that coating of plastic with ECM significantly enhances PD/passage (figure 1C) for all four ECMs tested in this work. The same was observed for CPD before reaching growth arrest as well (figure 1D). PD/passage and CPD before growth arrest for the ECMs were almost 2 times and 1.5 times higher than that for the non-ECM coated TCP respectively. However, out of the four ECMs tested, gelatin showed the best result in terms of PD/passage and final CPD before growth arrest. While cells on TCP had 2^8^ times increase in total cell number, cells on gelatin coated plastic had 2^14^ times increase in number before they reached growth arrest. However, it should be noted that the cells entered growth arrest at the same time irrespective of coating indicating no effect of the ECM on the culture duration at which growth arrest happens.

### ECM protein coating influences cell spread area for early and mid-passages

It is known that hMSCs increase their spread area with passage. Hence, to check the effect of ECM coating on the same, cell area for each condition was carefully measured before trypsinization during each passage. Figure 2 presents cell spread area from three passages: P3 (early passage), P6 (mid passage), P9 (late or growth arrest passage). It can be observed that average cell area and heterogeneity, as indicated by spread of the plot, increases with passage for all five conditions. However, as compared to other conditions, cells on non-ECM coated TCP start to spread more, even for early passages (figure 2AI). The average cell area at P3 was less than 4000 µm^2^ for all ECM proteins but was as high as 7143 ± 680 µm^2^ for non-ECM coated TCP. The cells were more heterogeneous on non-ECM coated TCP as well when compared to the ones on ECM proteins (figure 2AII and III).

**Figure 2:**
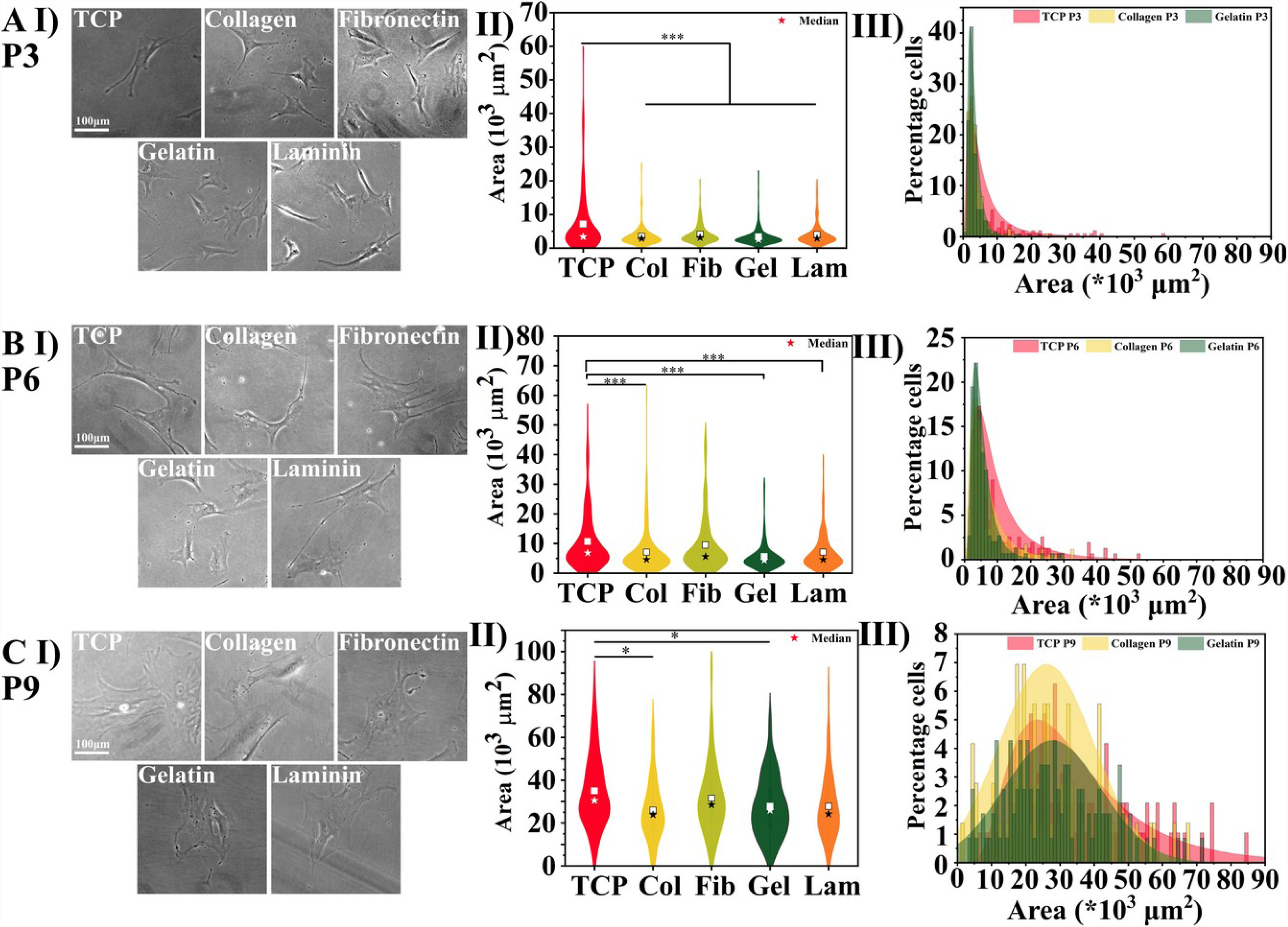
The figure shows the representative phase contrast images of BM-hMSCs seeded on ECM protein coated TCP across different passage A) P3 (N=3, n>170), B) P6 (N=3, n>140), C) P9 (N=3, n>65), and their respective cell spread area graphs (second column), followed by size histogram for TCP, collagen coated, and gelatin coated TCP. Filled black star represents the median and open square represents average. *P-*value *=p<0.05, **=p<0.01, ***p<0.001. Scale Bar 100 μm.

BM-hMSCs continued to increase their spreading across all conditions with passages, albeit at different rates. As the passage number increased from P3 to P6, the difference between cell area on non-ECM coated TCP and fibronectin became statistically insignificant, the averages being 10632±790 μm^2^ and 9479±743 μm^2^ respectively. However, the cells seeded on collagen, gelatin, and laminin coated TCP remained significantly smaller with areas 7087±570 μm^2^, 5593±379 μm^2^, and 7063±505 μm^2^ respectively, with gelatin being the lowest. Heterogeneity in cell spreading also started to increase, particularly for non-ECM coated TCP, as can be seen by comparing figure 2AIII and BIII. Many large cells started to appear on the non-ECM coated TCP substrate. While ∼60% of the cells on gelatin had area <5000 μm^2^, non-ECM coated TCP only had ∼35% of cells in that range.

As the cell culture further progressed to passage 9, cell spread area showed a tremendous increase as shown in the figure 2C. The cells cultured on non-ECM coated TCP showed an average area of 34988±1087 µm^2^. Though cells cultured on ECM protein coated TCP too had huge spreading (>25000 µm^2^, for all four ECM proteins), they were significantly less spread compared to their counterparts on non-ECM coated TCP. However, cells on all conditions started to show huge heterogeneity, as can be seen from figure 2C III.

To further investigate the fold change in cell spread area across different passages, we normalized the spread area with respect to non-ECM coated TCP P3 and the P3 of individual ECMs (figure S3). The area of the cells cultured on non-ECM coated TCP showed the highest fold change at P6 and P9, when normalized to the values at non-ECM coated TCP P3 (figure S3 A). On the other hand, when normalized to P3 of the respective ECM proteins, the highest fold change was observed for fibronectin at P6 and gelatin at P9 (figure S3).

### Cells cultured on gelatin coated TCP retained cellular stiffness

As the cellular spreading and senescence are related to cellular stiffness [10–12], we measured cellular stiffness at passages P5 and P10 across different coating conditions to estimate their role towards passage-related cell stiffness (figure 3). First, we noted that even at P5, cells on non-ECM coated TCP had much higher cellular stiffness compared to all other conditions. At P5, cells on collagen, laminin, and gelatin had very similar cell stiffnesses, which in turn was significantly lower than cells on non-ECM coated TCP. Cells on fibronectin had even lower rigidity than their counterparts on other ECMs. Further, when measured at P10, collagen, laminin, and fibronectin showed significant increase. In particular, the increase in cell stiffness on collagen and fibronectin was dramatic. Interestingly, non-ECM coated TCP and gelatin did not show any significant increase in cell stiffness, the cells on non-ECM coated TCP being approximately twice as stiff as the cells on gelatin at both passages, P5 and P10.

**Figure 3:**
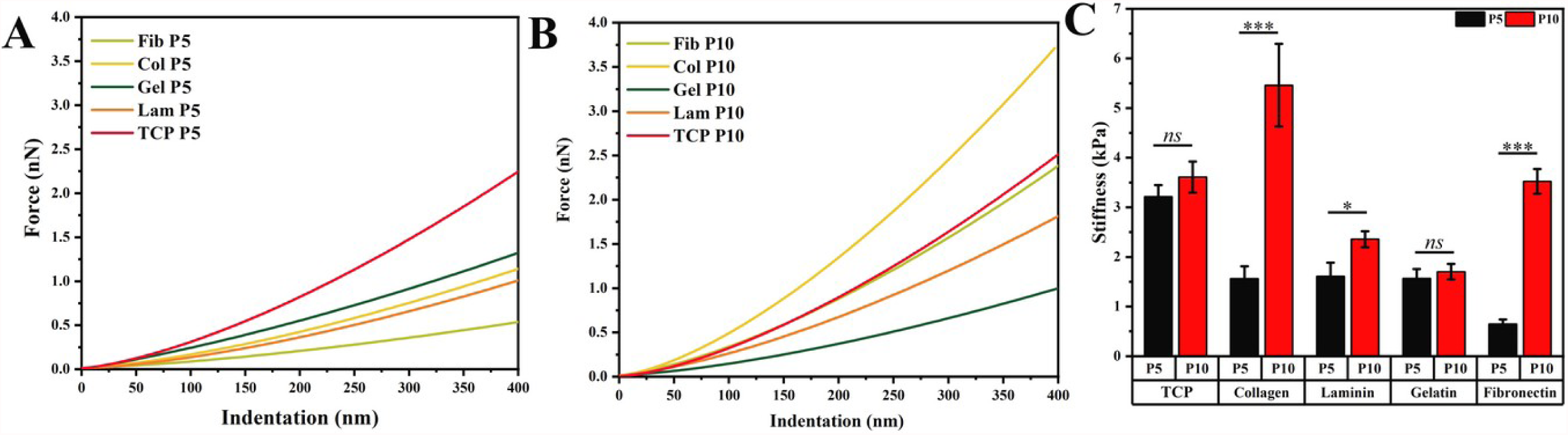
The line plots show the representative indentation curves of cells originating from different ECM-protein coated and non-coated TCP for passages: A) P5, and B) P10. C) Bar graph shows the average stiffness of cells originating from different ECM protein coated and non-coated TCP at passage P5 and P10. 25<n>76, mean±sem, P-value ***=p<0.001.

### Cells cultured on gelatin coated TCP retained differentiation potential

Multi-lineage differentiation potential is an important property of hMSCs which they lose on long term culture on plastic [13,14]. Hence, to check if gelatin coated TCP helped in retaining the cellular differentiation potential, we treated the cells from passage 4 (P4) and passage 8 (P8) with osteogenic and adipogenic induction media. After 14 days of induction the cells were stained with Oil Red O for adipogenic differentiation (top row of figure 4). After 21 days of induction cells were stained with Alizarin Red S for osteogenic differentiation (bottom row of figure 4). The cells at passage P8 retained their adipogenic and osteogenic differentiation.

**Figure 4:**
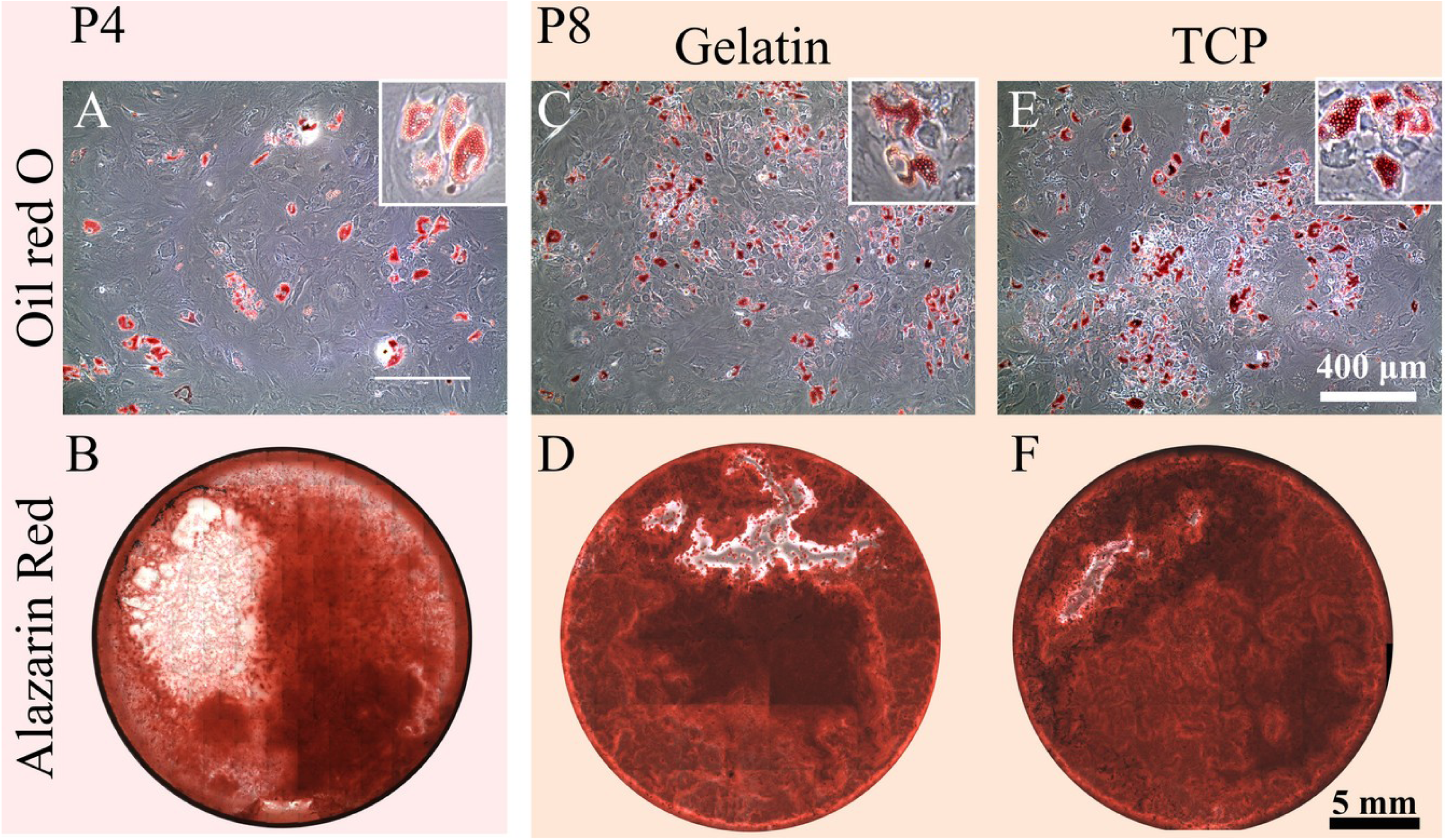
The first column of the figure shows the adipogenic differentiation (image A, cells stained with Oil Red O) and osteogenic differentiation (image B, cells stained with Alizarin Red S) of passage 4 (P4) cells. The second column shows the adipogenic differentiation (image C, cells stained with Oil Red O) and osteogenic differentiation (image D, cells stained with Alizarin Red S) of passage 8 (P8) cells expanded on gelatin coated TCP and the third column (Images E & F) shows differentiation of cells at P8 expanded on non-ECM coated TCP.

## Discussion

In this paper, we have investigated the effect of four different ECM proteins and non-ECM coated TCP on the growth, area, and stiffness of BM-hMSCs till they reach growth arrest. While various other papers earlier have studied the effect of surface coating on cell growth, the duration of study was for only a few hours to few days [1–4]. There has been no study to explore the effect of substrate coating on cell growth over extensive passaging. We have found that ECM protein coating indeed helped cell growth in terms of population doubling along with maintaining their differentiation potential. However, it had no effect on the duration that the cells needed to reach growth arrest which took place on all the ECMs around P9 which was after 42 days in culture (Fig 1B).

Studies using different cell types have demonstrated that the effect of natural ECM and peptide mimicking ECM proteins on cell proliferation is cell type dependent [2,4,15]. For example, using ECM protein mimetics, fibronectin was shown to have the greatest positive effect on the proliferation of human chorionic mesenchymal stem cells [4]. While for rat islet cells, Laminin was found to be the best ECM for cell proliferation [15]; for porcine lens epithelial cells, laminin coating did not have any effect better than regular tissue culture plates [16]. For umbilical cord derived haematopoietic progenitor cells, the effect of ECMs on cell proliferation was found either non-existent or limited [2].

An exclusive aspect of our work compared with the existing ones is the duration of culture. In none of the existing literature, the effect of ECM proteins was investigated over multiple passages till the point of growth arrest. Our data shows that for BM-hMSCs, although ECM coating improves the rate of population doubling, it does not have any effect on delaying the onset of growth arrest. In our earlier work, we have demonstrated that soft substrate can delay the onset of replicative senescence in BM-hMSCs [7], which was later supported by observations from other groups as well [16]. We demonstrated the same with patient derived keratinocytes as well [8]. However, ECM protein coating on TCP does not have any effect in delaying the senescence.

Another important parameter to study in this context is the cellular spreading. As reported earlier, cell spread area can be used as a measure of cell health. Cells become large and increasingly flattened when they enter senescence [7]. ECM coating on the substrate is known to influence cell spread area [17]. As cell passage increased, there was an increase in cell spread area which is in accordance with literature [7]. However, the fold change in cell spread area was minimal when cells were cultured on collagen and gelatin coated TCPs till P6. With increase in passage the cell spread area drastically increased and cells attained flattened morphology while entering growth arrest.

With increase in cell passage, literature has reported that cells show increased stiffness during expansion [18]. With our experiments, we found that at P5, cells on ECM coated plates had much lower stiffness compared to their counterparts on TCP. However, with increase in passage from P5 to P10, there was significant increase in cellular stiffness across all ECM proteins, particularly for collagen and fibronectin. Interestingly, stiffness of the cells on gelatin coated plates showed no increase in stiffness.

## Conclusion

In conclusion, we have demonstrated that coating the culture dish with ECM proteins can lead to higher cell number. This observation is important in the light of limited availability of BM-hMSCs for therapeutic use. Out of all the ECMs tested, we found the least expensive one, that is gelatin to be the best to support cell growth. Hence, gelatin coating on culture dish can facilitate higher cell number generation with minimal increment of financial burden. As the rate of population doubling (PD/passage) was highest on gelatin, it will also generate the required cell numbers the fastest. Our data demonstrates that it is advisable to check the effect of ECM before culturing any new cell type, particularly the ones with limited availability and growth.

## Author contribution

AM and PM conceptualized and designed the experiments, methodology. AM supervised the project. PM and SI performed all of the experiments. PM and SI did the analysis. PM prepared the figures. PM and AM interpreted the results. All three authors contributed to writing of the manuscript.

## Conflict of interest

Author declares no conflict of interest

## Acknowledgment

We thank IRCC, IIT Bombay for providing the Bio-AFM facility.

## Funding statement

This research was funded by Wadhwani Research Centre for Bioengineering (WRCB). We thank IRCC, IIT Bombay for providing the fellowship to PM, and WRCB for the salary of SI.

## SUPPLEMENTARY FIGURES

**Figure S1:**
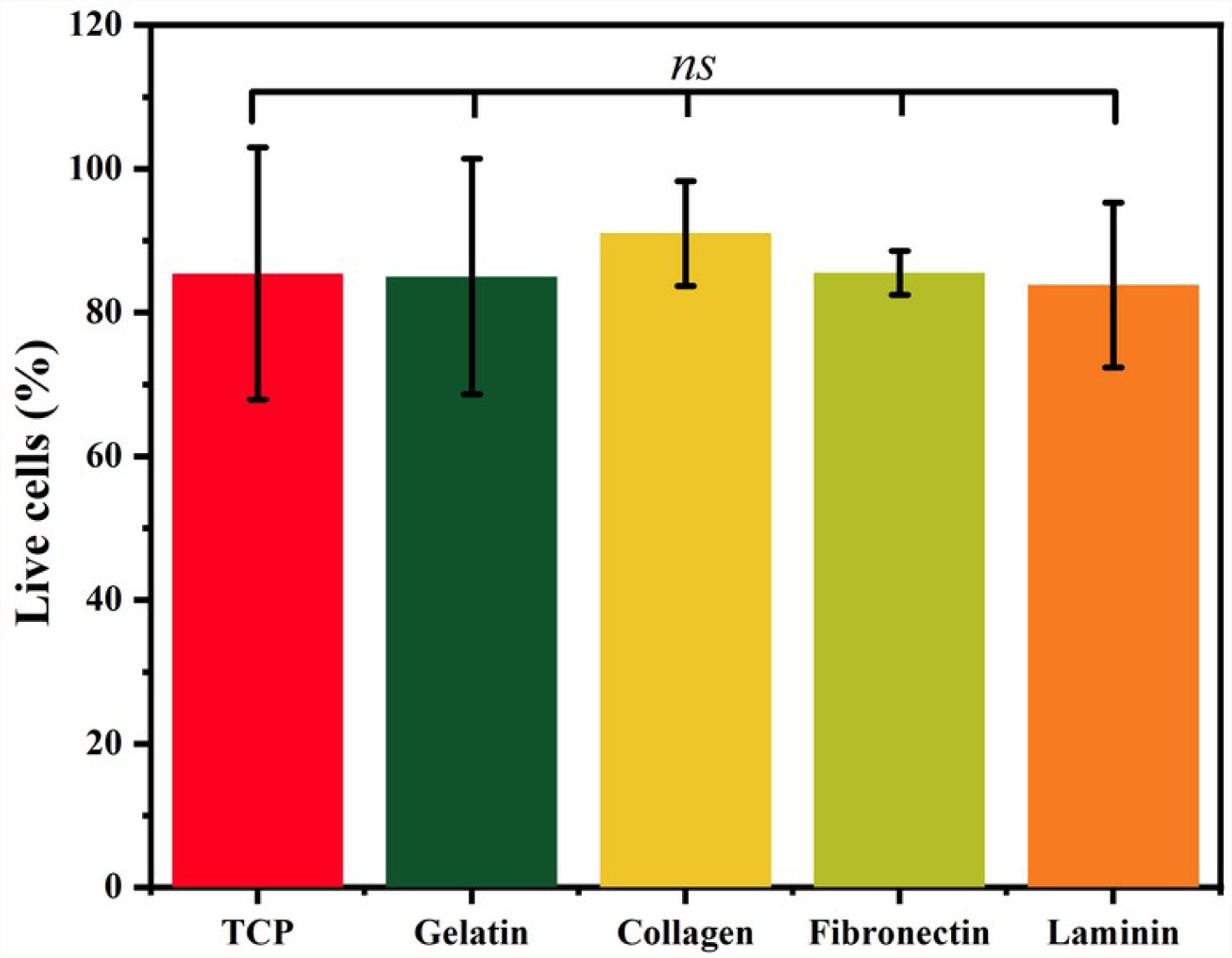
The bar graphs show the percentage of live cells after 24 hours of seeding on ECM protein-coated TCP and non-coated TCP. N=3, n>200.

**Figure S2:**
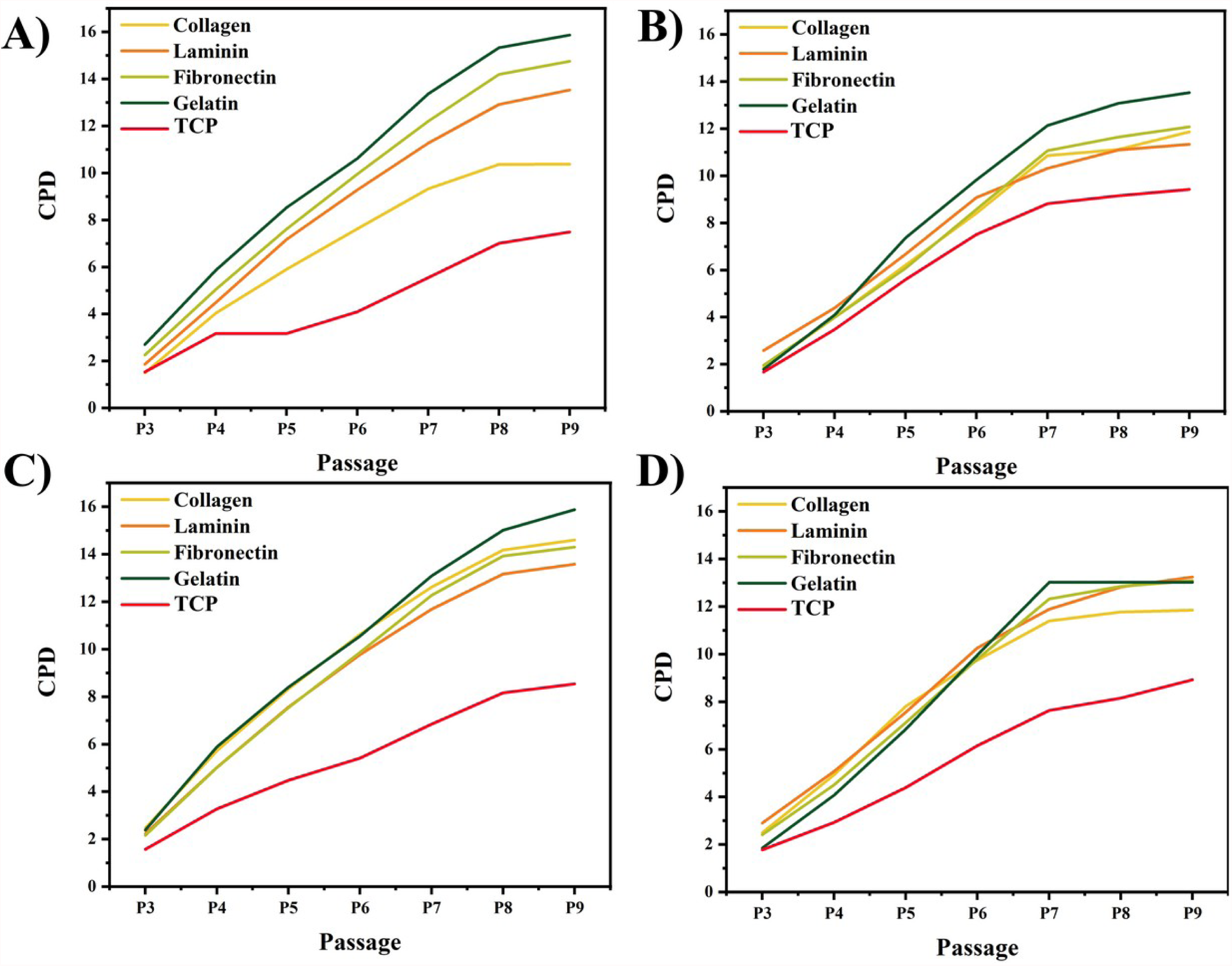
The line graphs show the CPD values for cells cultured on different ECM protein coated surfaces for four independent experiments.

**Figure S3:**
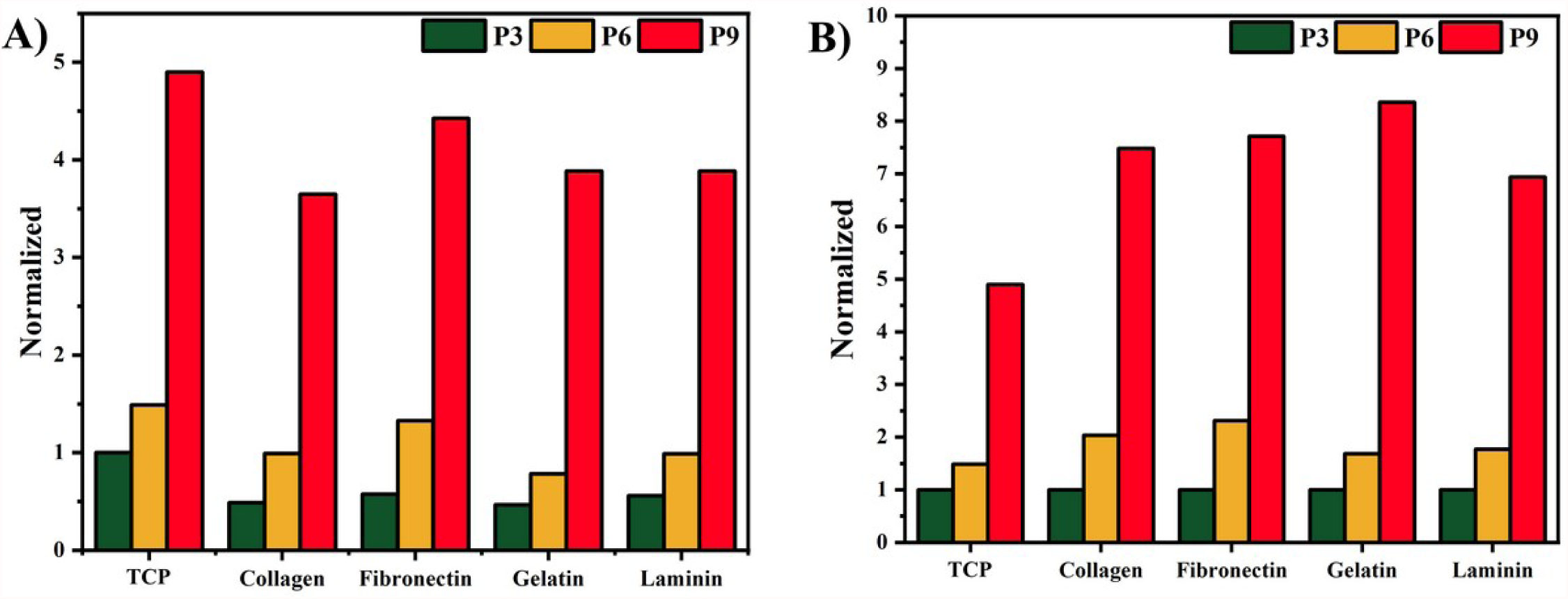
Bar graph shows the fold change in cell spread area when normalized to A) P3 cell spread area of non-ECM coated TCP and B) Cell spread area of the P3 of the respective coating.

## Notes

### Competing Interest Statement

The authors have declared no competing interest.

